# Rewiring of the promoter-enhancer interactome and regulatory landscape in glioblastoma orchestrates gene expression underlying neurogliomal synaptic communication

**DOI:** 10.1101/2022.11.16.516797

**Authors:** Chaitali Chakraborty, Itzel Nissen, Craig A. Vincent, Anna-Carin Hägglund, Andreas Hörnblad, Silvia Remeseiro

## Abstract

Chromatin organization controls transcription by modulating 3D-interactions between enhancers and promoters in the nucleus. Alterations in epigenetic states and 3D-chromatin organization result in gene expression changes contributing to cancer pathogenesis. Here, we mapped the promoter-enhancer interactome and regulatory landscape of glioblastoma, the most aggressive primary brain tumour. Our data reveals profound rewiring of promoter-enhancer interactions, chromatin accessibility and redistribution of histone marks across the four glioblastoma subtypes. This leads to loss of long-range regulatory interactions and overall activation of promoters, which orchestrate changes in the expression of genes associated to glutamatergic synapses, axon guidance, axonogenesis and chromatin remodeling. SMAD3 and PITX1 emerge as the major transcription factors controlling genes related to synapse organization and axon guidance. Inhibition of SMAD3 and neuronal activity stimulation cooperate to promote cell proliferation of glioblastoma cells in co-culture with glutamatergic neurons. Our findings provide mechanistic insight into the regulatory networks that mediate neurogliomal synaptic communication.

## Introduction

Gene regulation critically relies on regulatory sequences such as enhancers, which control the spatial and temporal specificity of gene expression. In vertebrates, enhancers are often located hundreds of kilobases (kb) to even megabases (Mb) away from their target gene promoters. This long-range action of enhancers over gene promoters is facilitated via the structural organization of the genome in the 3D nuclear space, mainly within sub-megabase domains known as TADs (Topologically Associating Domains)^1,2^, which emerge from multiple nested loops by loop extrusion mechanisms^3^. This topological organization of the chromatin allows for physical proximity between enhancers and promoters, constraining the range of action of enhancers and therefore setting the stage for the specificity of promoter-enhancer interactions. Enhancer activity is intimately linked to epigenetic status and, thus, enhancers display different states (i.e., active, silenced, primed or poised) depending on the combination of histone marks and other chromatin features^4–8^. Genome-wide enhancer maps have linked risk variants to disease genes^9^, and most genomic variants that predispose to cancer are located in non-coding regions with potential to act as *cis*-regulatory elements^10^. Various evidence supports the role of alterations in the regulatory and topological landscape in different types of cancer. Aberrant super-enhancer function provides oncogenic properties, and cancer cells can acquire super-enhancers to drive expression of oncogenes^11–13^. Proto-oncogenes can become activated by disruption of chromosome insulated neighbourhoods^14^, and systematic occurrence of structural rearrangements in *cis*-regulatory elements (e.g., enhancer hijacking) mediates dysregulation in cancer^15,16^.

Glioblastoma (GB) (WHO grade IV astrocytoma) is the most malignant form in the wide spectrum of gliomas, the most common primary brain tumours. With a 5-year survival rate of only 3-4%^17^, glioblastoma prognosis has not improved considerably in the last decades. Glioblastomas develop rapidly and manifest after a short clinical history of usually less than 3 months^18^. No uniform aetiology has been identified, and mechanistic understanding of GB initiation and progression is difficult given the complexity of genomic, epigenomic, metabolic and microenvironment events contributing to disease. Extensive inter- and intra-tumour heterogeneity are characteristics of glioblastoma that have complicated the finding of specific and targeted therapies.

Epigenetic alterations play a central role in the aetiology of gliomas and are used for molecular classification^19^. In higher-grade gliomas, diverse alterations in genes encoding chromatin remodelers and epigenetics-related enzymes have been described in conjunction with deregulation of the epigenetic landscape and subsequent gene expression alterations^20–22^. Early studies identified four glioblastoma subtypes defined by aberrations and gene expression changes in *EGFR*, *NF1*, and *PDGFRA/IDH1*, corresponding to classical, mesenchymal and neural/proneural subtypes, respectively^23^. Single-cell studies have recently identified that glioblastoma cells display plasticity and transition between four main cellular states influenced by the tumour microenvironment^24^. In the last few years, researchers have begun to integrate transcriptomic analysis with chromatin and epigenetic profiles, DNA methylomes or chromatin architecture data in glioblastomas^25–30^, evidencing that glioblastoma is an entity distinguishable from lower grade gliomas. However, despite the efforts to further classify glioblastoma in molecular subtypes and to identify relevant subpopulations, this has not yet been translated into a clinical benefit.

In this study, we apply a multi-omics approach to map the promoter-enhancer interactome and the regulatory landscape of glioblastoma, including histone marks and chromatin accessibility, in a panel of 15 patient-derived glioblastoma cell lines representing all four expression subtypes and normal human astrocytes as control. We observed a rewiring of the promoter-enhancer interactions, changes in chromatin accessibility and a redistribution of histone marks across all four expression subtypes in glioblastoma. These changes in the regulatory and topological landscapes lead to a significant loss of long-range regulatory interactions and an overall activation of promoter-hubs. This orchestrates changes in the expression of genes associated to synapses, in particular glutamatergic synapses, as well as axon guidance and axonogenesis, and chromatin binding/remodeling. Motif search analysis reveals the transcription factors (TFs) SMAD3 and PITX1 as major direct regulators of a set of downstream target genes related to synaptic contacts and axon guidance. In addition, we functionally demonstrate that inhibition of SMAD3 and stimulation of neural activity additively cooperate to promote proliferation of glioblastoma cells. After the recent discovery of neurogliomal synapses^31–33^, that drive tumour progression by relaying neuronal activity to tumour cells, our data offers novel mechanistic insight into the gene regulatory networks that mediate the neurogliomal synaptic communication in glioblastoma.

## Results

### A map of the regulatory landscape and promoter-enhancer interactome in glioblastoma

To obtain a map of the enhancer landscape and promoter-enhancer interactome in glioblastoma, we performed a multi-omics approach in a panel of 15 patient-derived glioblastoma cell lines alongside normal human astrocytes as a control (Fig. 1a). The 15 patient-derived glioblastoma cell lines were obtained through the human glioblastoma cell culture (HGCC) resource^34^, a panel of newly established and well characterized glioblastoma lines derived from GB patient surgical samples, that represent all four expression subtypes (i.e., classical, mesenchymal, neural and proneural). Normal human astrocytes were selected as control since astrocytes are considered to be a cell of origin for glioblastoma^35,36^.

**Figure 1:**
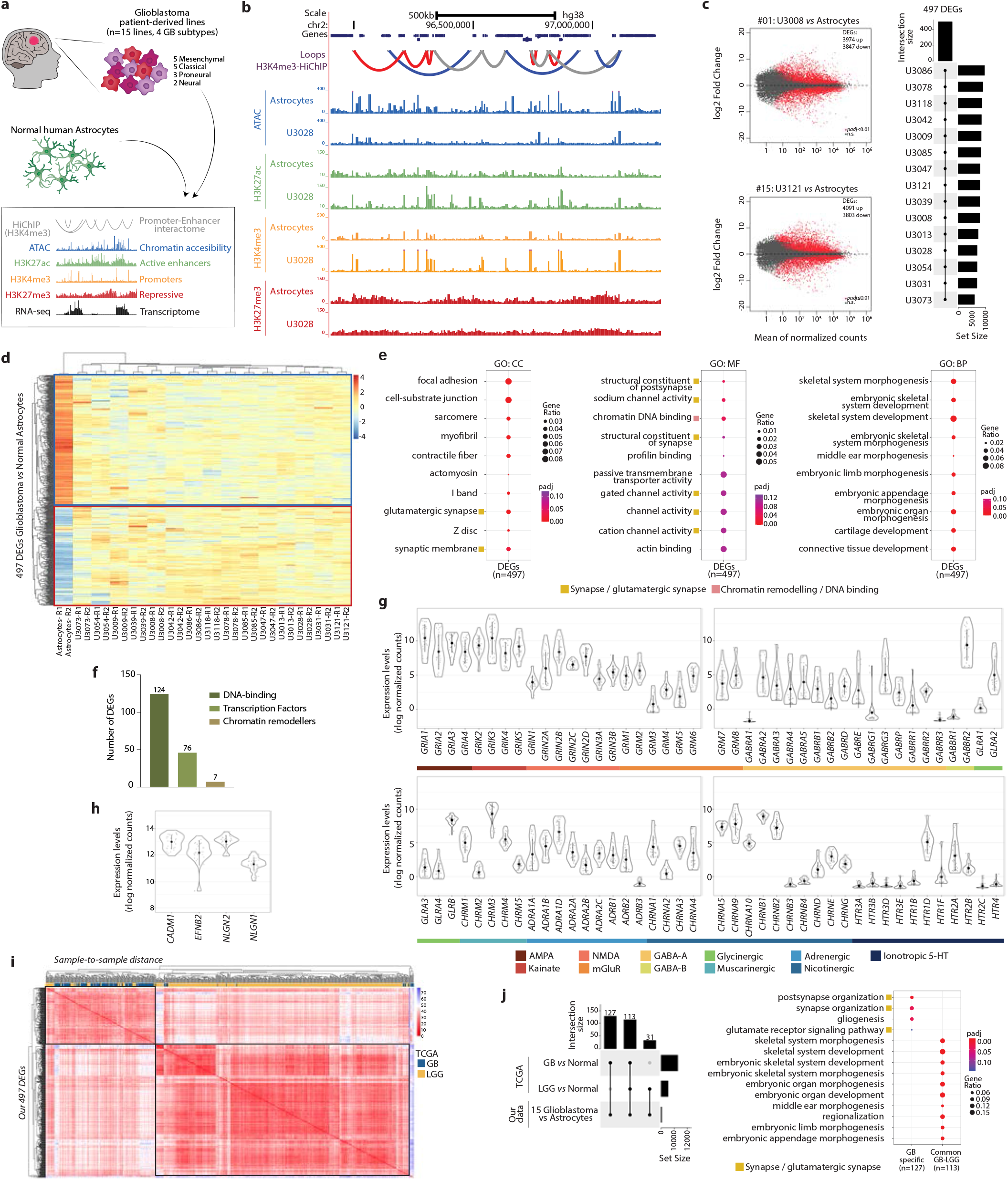
Gene expression changes common to all four glioblastoma subtypes relate to synapse organization, glutamatergic synapse and chromatin binding. a) Experimental workflow. b) Genomic distribution of HiChIP loops, chromatin accessible regions by ATAC, and H3K27ac, H3K4me3 and H3K27me3 peaks in normal astrocytes and one of the 15 glioblastoma cell lines (U3028). c) Differentially expressed genes (DEGs, red dots) in two representative glioblastoma lines *vs* normal astrocytes (left) and intersection of the 15 pairwise differential expression analysis resulting in 497 DEGs (right). d) Expression of 497 DEGs in all 15 glioblastoma lines and normal astrocytes as determined by RNA-seq (rlog normalized counts). e) Top 10 Gene Ontology terms enriched in the 497 DEGs. f) Number of DEGs annotated as DNA-binding, transcription factors or chromatin remodelers. g-h) Expression of neurotransmitter receptor genes (g) and synaptogenesis markers (h) in the 15 glioblastoma lines. i) Sample-to-sample distance for TCGA samples annotated as GB (glioblastoma, n=156) or LGG (lower grade glioma, n=511). j) Intersections between our 497 DEGs and the differentially expressed genes from TCGA tumour samples, either GB *vs* normal or LGG *vs* normal tissue (left). Top 10 GO terms enriched in the 127 GB-specific genes or 113 genes common to GB and LGG (right).

We mapped the promoter-enhancer interactome by HiChIP^37^, a protein-centric chromatin conformation capture method, using an antibody specific to the promoter mark H3K4me3. Assay for Transposase-Accessible Chromatin with sequencing (ATAC-seq)^38^ was used to map chromatin accessibility genome-wide. Via ChIP-seq (Chromatin ImmunoPrecipitation-sequencing) we determined binding sites for histone modifications: the active enhancer mark H3K27ac, the repressive mark H3K27me3, and H3K4me3 predominantly associated to gene promoters. Transcriptome profiling of each line was done by RNA-seq. Our data constitutes a comprehensive map of the regulatory landscape in glioblastoma across all four subtypes, and provides functionally relevant topological information by mapping of the promoter-enhancer interactome (Fig. 1b). In the next sections, we compare the regulatory and topological landscape of glioblastoma to the control astrocytes, while focusing on the features present in all four GB subtypes.

### Gene expression changes across all four GB subtypes are associated to synapse organization, glutamatergic synapses and chromatin binding

Transcriptome profiles were obtained for each of the 15 patient-derived cell lines and the normal human astrocytes by bulk RNA-seq. To identify gene expression changes across all four GB subtypes, we first performed a pair-wise differential expression analysis of each glioblastoma line *vs* the control astrocytes (Fig. 1c and Supplementary Fig. 1a), and then intersected the differentially expressed genes (DEGs) resulting from the 15 pair-wise comparisons (Fig. 1c). We thus identified a set of 497 DEGs that are differentially expressed across all four GB subtypes (Fig. 1d, Supplementary Table S1). Gene Ontology (GO) analysis of this gene set revealed a significant enrichment for terms associated to synapses, and in particular glutamatergic synapses, channel activity and chromatin DNA binding (Fig. 1e). Other GO terms enriched were those related to morphogenesis and development, processes well known to be dysregulated in cancer. A closer analysis of the 497 DEGs showed that 124 genes are annotated as DNA binding, 76 genes encode for Transcription Factors (TFs) and additional 7 genes encode chromatin remodelers (Fig. 1f), evidencing common changes in chromatin-related genes and TFs across the four GB subtypes.

In addition, we detected expression of neurotransmitter receptor genes in all 15 glioblastoma lines (Fig. 1g), including expression of AMPA, kainate, NMDA and mGluR, which are receptors that respond to glutamate, and in accordance with the glutamatergic identity of the neurogliomal synapses^31^. In particular, the kainate receptor gene *GRIK2* and the NMDA receptor gene *GRIN2C* are also differentially expressed across all four GB subtypes in comparison to normal astrocytes. The expression of four known synaptogenesis markers^39^ is also detected in our panel of glioblastoma lines (Fig. 1h), in agreement with the emergence of cell subpopulations with synaptogenic properties during glioma progression^40^. Notably, high expression level of the synaptogenesis marker *EFNB2* correlates with lower overall survival of GB patients (TCGA data, Supplementary Fig. 1b), evidencing its clinical relevance in glioblastoma.

Importantly, the 497 DEGs dysregulated across all four subtypes also constitute a gene signature that segregates glioblastoma (GB) from lower grade gliomas (LGG). In a panel of more than 600 tumour samples available at The Cancer Genome Atlas (TCGA), including glioblastoma tumours (GB, n=156) and lower grade gliomas (LGG, n=511), the expression of our 497 DEGs can accurately segregate GB from LGG (Fig. 1i). We then performed a differential expression analysis of the TCGA RNA-seq data from either GB or LGG samples with respect to the corresponding controls, followed by intersection with the 497 DEGs from our data. As a result, we obtained a subset of 127 GB-specific genes and 113 genes “common to all gliomas”, the latter resulting from the intersection of both GB gene sets and the LGG gene set (Fig. 1j). Interestingly, GO analysis demonstrated that the 113 genes that are common to all gliomas are preferentially related to morphogenetic and developmental processes, whereas the subset of 127 GB-specific genes is very specifically associated to synapse organization, gliogenesis and glutamate receptor signalling (Fig. 1j). The neurogliomal synapses (i.e., bona fide synapses between presynaptic neurons and postsynaptic glioma cells) utilize glutamate receptors and trigger postsynaptic signals, which in turn affect proliferation and migration of the tumour cells^31–33^. Our analysis reveals that gene expression changes affecting synapses, in particular glutamate receptor signalling, are associated more strongly with glioblastoma samples when compared with lower grade gliomas. This points to certain genes that may be predominant regulators of neural function in glioblastoma pathogenesis.

### Loss of long-range regulatory interactions and gain of promoter hub interactions characterise the 3D organisation of the glioblastoma genome

Control of transcription is exerted by the physical interaction between enhancers and promoters through a non-linear relationship^41^. To chart these physical interactions genome-wide in glioblastoma, we performed HiChIP with an antibody against the promoter and enhancer mark H3K4me3. We thus obtained a map of the promoter interactome in our panel of glioblastoma lines representing all four subtypes, including not only promoter-enhancer (P-E) but also promoter-promoter (P-P) and enhancer-enhancer (E-E) interactions (Fig. 2a). A comparison of the loops detected by H3K4me3-HiChIP shows that 4316 loops detected in the normal astrocytes are also present in the GB samples, while an additional 5633 are differential loops: 2125 loops are lost (i.e., astrocyte-specific) and 3508 loops are gained in glioblastoma (i.e., GB-specific) (Fig. 2b). Interestingly, 86.6% of the lost loops involve enhancer interactions (P-E and E-E), while 85.4% of the gained loops involve exclusively promoter-promoter (P-P) interactions (Fig. 2c). Analysis of only multi-anchor loops (i.e., more than two anchor sites, 78% of total loops) shows that a major fraction of these are gained multi-anchor P-P loops (87.4%, Supplementary Fig. 2a), suggesting that gained P-P interactions in glioblastoma occur mainly in promoter hubs. Importantly, the length of the lost loops (median ~254kb) is significantly higher than that of the gained loops (median ~114kb, p-value=1.36e^−290^), supporting a preferential loss of long-range interactions in glioblastoma (Fig. 2d). These changes in the promoter-enhancer interactome are accompanied by gene expression changes: 183 genes located at the anchors of differential loops are differentially expressed across the 15 patient-derived glioblastoma lines (Supplementary Fig. 2b), out of which 55 (29.7%) encode for transcription factors, chromatin remodelers and other DNA-binding proteins (Fig. 2e). Our map of the enhancer-promoter interactome reveals topological changes that include loss of long-range regulatory interactions and gain of promoter hub interactions in all four glioblastoma subtypes.

**Figure 2:**
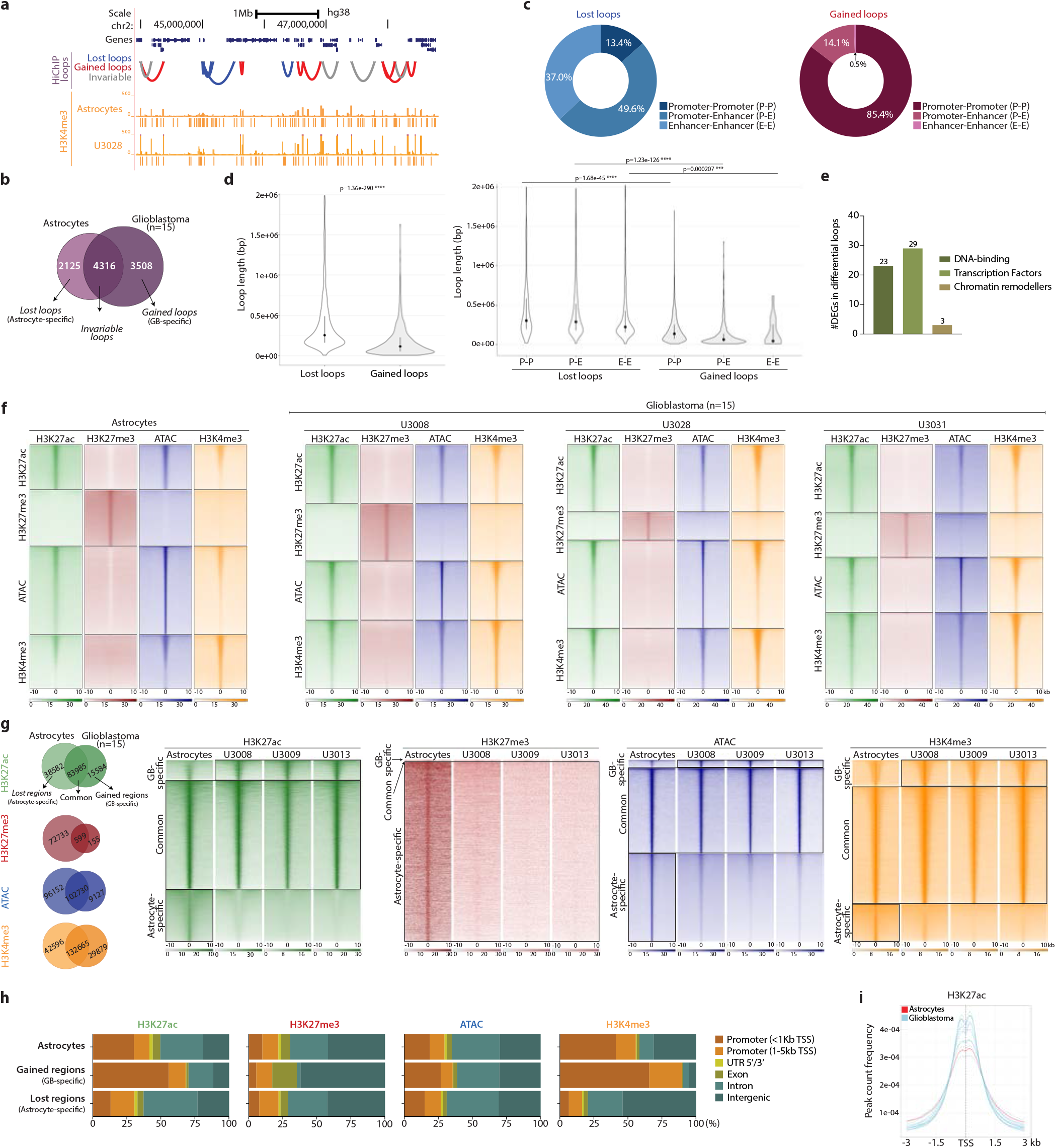
Loss of long-range regulatory interactions and changes in the regulatory landscape in glioblastoma. a) Distribution of HiChIP loops and H3K4me3 peaks in a region of chromosome 2. b) Intersection of HiChIP loops in the 15 glioblastoma lines and normal astrocytes defines lost loops (“astrocyte-specific”) and gained loops (“GB-specific”) in glioblastoma. c) Annotation of lost and gained loops as P-P, P-E or E-E loops. d) Length (bp) of lost and gained loops (t-test, Benjamini correction). e) Number of DEGs at the anchors of differential loops annotated as DNA-binding, transcription factors or chromatin remodelers. f) Read distribution of histone marks and ATAC around the indicated ±10kb regions in astrocytes and three representative glioblastoma lines. g) Redistribution of histone marks and ATAC regions in glioblastoma. h) Genomic annotation of differential histone peaks and ATAC regions in glioblastoma *vs* normal astrocytes. i) H3K27ac peak count frequency around the TSSs.

### Remodeling of the regulatory landscape in glioblastoma is characterized by loss of regulatory elements and activation of promoters

Depending on the combination of histone marks and other chromatin features, enhancers can present different states from active to silenced, primed or poised^4–8^. To map the regulatory landscape in glioblastoma, we profiled by ChIP-seq the binding of H3K27ac (active enhancers), H3K27me3 (polycomb-repressed) and H3K4me3 (promoters and enhancers), together with chromatin accessibility by ATAC-seq (Fig. 2f, Supplementary Table S2). Multiinter intersection of the binding sites in the 15 glioblastoma lines *vs* the control astrocytes, reveals a redistribution of histone marks and changes in chromatin accessibility occurring across the four GB subtypes (Fig. 2g). This is evidenced by the loss of binding sites that were present in astrocytes (i.e., “lost regions”) and the remobilization of histone marks to new genomic positions (i.e., “gained regions”) in glioblastoma. Similarly, a fraction of the chromatin accessible regions defined by ATAC are altered in glioblastoma. Analysis of genomic annotation of the differential regions shows that the remobilization of histone marks results in a loss of active marks at distant elements and an accumulation at gene promoters, while repressive marks are lost from intergenic regions (Fig. 2h,i).

As part of the integration of our multi-omics data genome-wide (Fig. 3a), we used ChromHMM to integrate our datasets and we characterized eight different chromatin states in the normal human astrocytes (Fig. 3b). Plotting the signal of the histone marks and ATAC-seq around the astrocytes’ chromatin states displayed clear differences between the glioblastoma lines and the control astrocytes (Fig. 3c-e). H3K27ac signal increases in poised and weak promoters but decreases around both strong and weak enhancers (Fig. 3d), further supporting the activation of poised/weak promoters and the loss of regulatory activity at enhancers. In addition, the repressive mark H3K27me3 decreases at poised promoters and polycomb-repressed states in glioblastoma (Fig. 3e). These chromatin changes in glioblastoma are also accompanied by increased expression levels at inactive/poised promoters and in the proximity of polycomb-repressed regions (Fig. 3f). Such rewiring of the regulatory landscape, together with the changes in the promoter-enhancer interactome and gene expression levels, supports a loss of long-range regulatory interactions and overall activation of hub promoters across all four GB subtypes.

**Figure 3:**
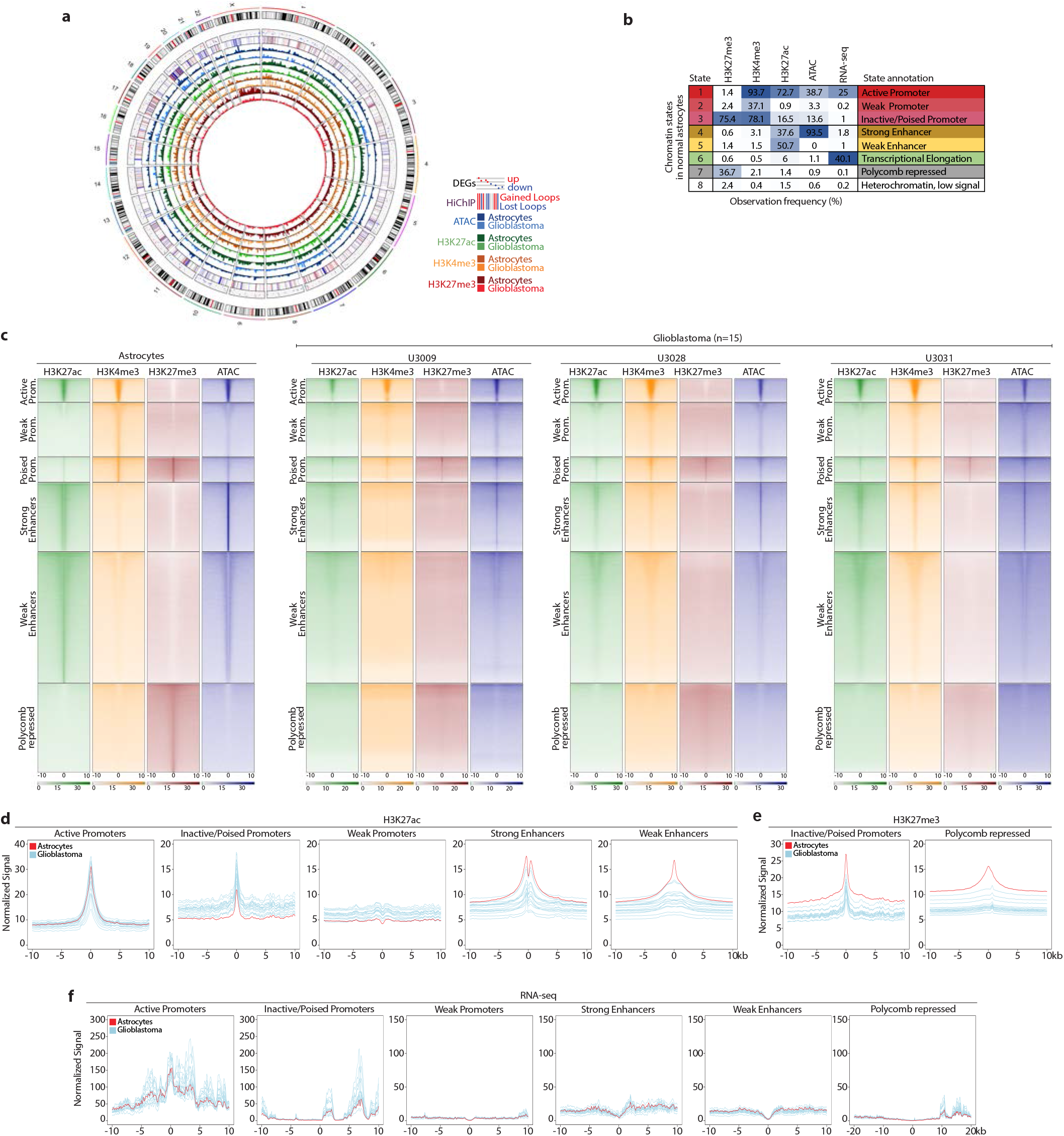
Remodeling of the regulatory landscape in glioblastoma includes reduction of active marks at enhancers and activation of promoters. a) Genome-wide integration of all the multi-omics data. b) Chromatin states defined by chromHMM in normal astrocytes. c) Read distribution of histone marks and ATAC across the various chromatin states in astrocytes and three representative glioblastoma lines. d-f) Density plots representing the normalized H3K27ac (d), H3K27me3 (e) and RNA-seq (f) signal around the astrocytes’ chromatin states in glioblastoma (blue) *vs* astrocytes (red).

### Remobilization of active chromatin marks around genes associated to glutamatergic synapses, axon guidance and chromatin remodeling

The rewiring in the enhancer landscape in glioblastoma leads to an increased chromatin accessibility and an accumulation of active histone marks at gene promoters. Gene Ontology analysis revealed that those genes near the newly occupied active positions (i.e., gained H3K27ac and ATAC regions) are associated to glutamatergic synapses, axon guidance and axonogenesis, as well as chromatin remodelling/DNA binding (Fig. 4a). Importantly, a major fraction of the genes associated to these biological processes are differentially expressed across the four glioblastoma subtypes (Fig. 4b and Supplementary Fig. 3a). Moreover, the loss of repression (i.e., lost H3K27me3 regions) occurs in the vicinity of genes related to calcium ion homeostasis, ion transport and synaptic transmission (Supplementary Fig. 3b). We therefore observed an accumulation of active marks around genes related to synapses, axon guidance and axonogenesis, and loss of repression of other genes associated to ion transport and synaptic transmission. Altogether, this suggests that the rewiring of the regulatory landscape in glioblastoma orchestrates a series of gene expression changes that contribute to the synaptic communication between neurons and glioma cells.

**Figure 4:**
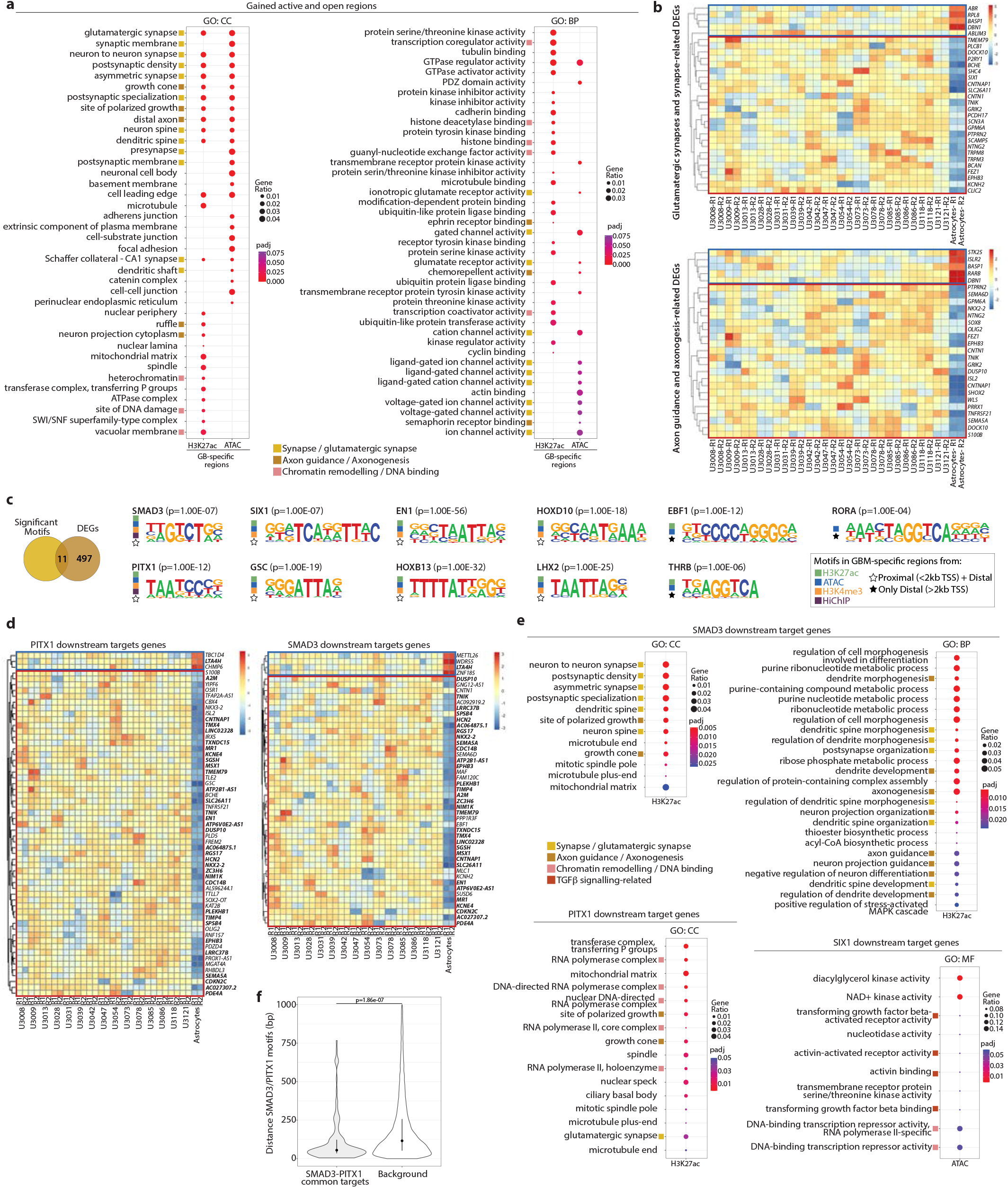
SMAD3 and PITX1 regulatory networks control genes associated to (glutamatergic) synapse organization and axonogenesis in glioblastoma. a) Top 25 GO terms enriched for genes proximal to gained active and open regions in glioblastoma (peaks <2kb TSS). b) Expression of DEGs related to synapse and glutamatergic synapse (top), and axon guidance and axonogenesis (bottom) located <2kb from a gained active/open peak. c) Intersection of the 497 DEGs with the HOMER significantly enriched motifs at the gained regions or anchors of differential loops identifies 11 key TFs. d) Expression of PITX1 and SMAD3 downstream target genes (i.e., motif enrichment at a differential peak <2kb TSS) among the DEGs in glioblastoma *vs* normal astrocytes (common targets in bold). e) Top GO terms enriched for SMAD3, PITX1 and SIX1 proximal downstream target genes. f) Distance between SMAD3 and PITX1 motifs at the promoters of the 33 common SMAD3/PITX1 target genes (left, median=56bp) *vs* background model (right, median=124bp). (Wilcoxon test).

### SMAD3 and PITX1 regulatory networks control synapse organization and axonogenesis in glioblastoma

A search for transcription factor binding site (TFBS) motifs revealed the enrichment of 11 key TFBS motifs within the newly occupied active regions in glioblastoma (Fig. 4c, Supplementary Fig. 3c,d). Not only are the motifs of these 11 TFs enriched in the gained active and open regions, but the genes encoding these 11 TFs are also differentially expressed across all four GB subtypes (i.e., 11 TFs out of the 76 TFs differentially expressed, Fig. 1f). Moreover, a significant fraction of their direct downstream target genes (i.e., motif within a gained peak <2kb TSS) are also differentially expressed in the four GB subtypes (Fig. 4d, Supplementary Fig. 4a). These findings reflect the impact that chromatin mark redistribution and accessibility changes around these TFBS has on gene expression.

GO analysis of the downstream target genes of these 11 TFs points to SMAD3 and PITX1 as major players in the regulatory networks that mediate neurogliomal synaptic communication (Fig. 4e, Supplementary Fig. 4b,c). Various GO terms related to synapse density, postsynaptic organization, axon guidance and axonogenesis are significantly enriched in the case of SMAD3 downstream targets, and in particular glutamatergic synapses for the PITX1 downstream target genes (Fig. 4e). This is in agreement with the reported glutamatergic identity of the neurogliomal synapses^31^. GO terms related to TGFβ receptor activity are significantly enriched among the SIX1 downstream targets (Fig. 4e), which links TGFβ signalling via SMAD3 to the synaptic communication between neurons and glioma cells. Importantly, 70.2% of the SMAD3 target genes differentially expressed in glioblastoma (i.e., 33 out of 47) are also PITX1 downstream targets (Fig. 4d, in bold). The common SMAD3/PITX1 targets include genes encoding semaphorins and ephrins involved in axon guidance^42,43^ such as *SEMA5A* and *EPHB3*, the latter of which is also known to participate in the development of excitatory synapses^44^. Other SMAD3/PITX1 targets include the transcription factor *NKX2-2* that is involved in regulating axon guidance^45^; *HCN2* and *KCNE4* that encode for gated channels, the latter considered to regulate neurotransmitter release^46^; as well as *TNIK* which is implicated in glutamatergic signalling, where it binds to NMDA receptors and is required for AMPA expression and synaptic function^47^.

Notably, SMAD3 and PITX1 binding motifs are also located in close proximity (median distance=56bp, Fig. 4f) at the promoters of the 33 common target genes differentially expressed across the four glioblastoma subtypes. In addition, SMAD3 and PITX1 binding motifs are enriched at the anchors of the differential loops in glioblastoma (Supplementary Fig. 3e). Interestingly, Smad proteins have been previously reported to bind CTCF sites in a CTCF-dependent manner in flies^48^ and SMAD3 interacts with CTCF in mammalian cells^49^. Apart from SMAD3/PITX1, additional regulatory networks could contribute to a certain extent to the neural role in glioblastoma pathogenesis, since other transcription factors such as EBF1, THRB, EN1, HOXB13 and HOXD10 are also enriched at gained active regions distal to genes involved in axonogenesis and axon guidance (i.e., motif within a gained peak >2kb TSS) (Supplementary Fig. 4c, Supplementary Fig. 5a).

Comparison of *SMAD3-PITX1* and *SMAD3-SIX1* co-expression showed a significant negative correlation between their expression levels, both in our panel of 15 patient-derived samples (Fig. 5a) and in 667 TCGA glioma samples (Fig. 5b). While *SMAD3* is downregulated, *PITX1* and *SIX1* are upregulated in glioblastoma (Fig. 1d). Moreover, low *SMAD3* and high *PITX1* expression levels correlate with lower overall survival in patients (TCGA data, Fig. 5c), pointing to their clinical relevance. It is important to highlight that both the *SMAD3* and *PITX1* loci present topological and regulatory changes (i.e., differential loops, redistribution of chromatin marks and chromatin accessibility) in comparison to normal astrocytes (Fig. 5d). Also, both *SMAD3* and *PITX1* genes are differentially expressed in all four GB subtypes, and their respective TFBS motifs are enriched at the gained active/open regions at the promoters of genes related to synaptic function, axon guidance and axonogenesis. This altogether indicates the presence of a regulatory network involving SMAD3 and PITX1, and it suggests that they may be the most prominent TFs regulating the neurogliomal synaptic interaction and axonogenesis in glioblastoma.

**Figure 5:**
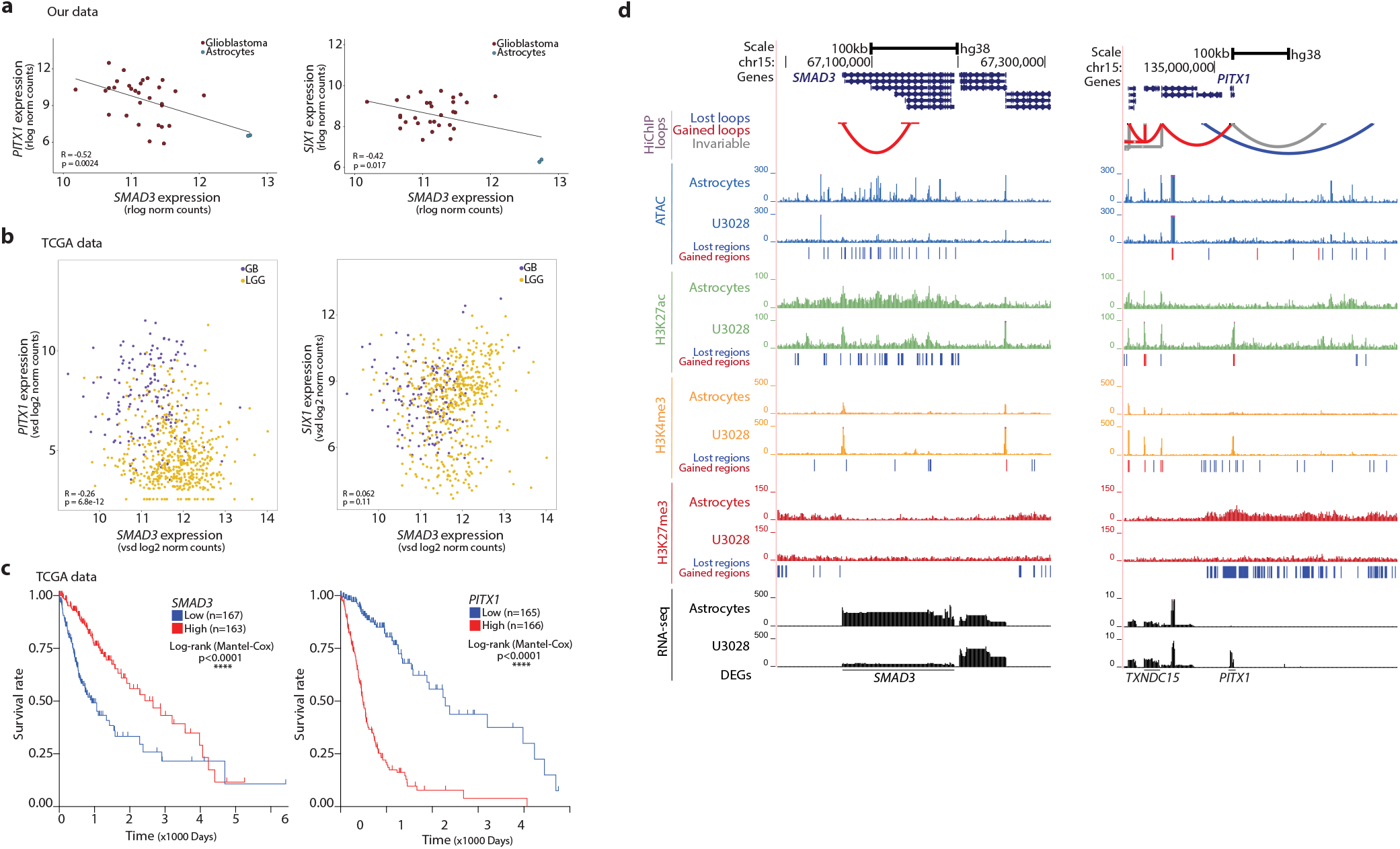
Regulatory alterations in the *SMAD3* and *PITX1* loci accompanied by inversely correlated gene expression and association to lower survival in patients. a-b) Scatterplots showing co-expression of SMAD3-PITX1 (left) or SMAD3-SIX1 (right) expressed as normalized counts in our data (a) and TCGA datasets (b) (Pearson correlations). c) Kaplan-Meier survival curves plotted for patients stratified by quartiles of SMAD3 or PITX1 expression (TCGA data) (log-rank Mantel-Cox test). d) Changes in the enhancer landscape and promoter-enhancer interactome at the SMAD3 and PITX1 loci (U3028 depicted as representative glioblastoma line).

### SMAD3 inhibition and neuronal activity stimulation cooperate to promote cell proliferation in glioblastoma

To functionally support the role of TGFβ signalling and in particular SMAD3 in this context, we tested the effect of their inhibition on the proliferation of glioblastoma cells. First, we induced the reprogramming of glutamatergic neurons (ab259259) from human iPSCs (induced pluripotent stem cells) (Fig. 6a-c), and then established co-cultures of glutamatergic neurons and the glioblastoma line U251-GFP at different time points (Fig. 6a,d). By live-cell imaging, we determined the proliferation of the glioblastoma cells in co-culture with glutamatergic neurons upon stimulation of neuronal activity alone or in combination with inhibition of either SMAD3 or the TGFβ receptor ALK5. Treatment with the SMAD3-specific inhibitor SIS3 significantly increases the proliferation of glioblastoma cells in co-culture with glutamatergic neurons, both at early and late time-points (days 6-9 and 14-17, respectively) (Fig. 6e,f). Stimulation of neuronal activity by picrotoxin has only modest effects on U251 proliferation at the highest doses, and only in more mature neurons (day 14-17) (Supplementary Fig. 6a). Importantly, combination of SMAD3 inhibition with stimulation of neuronal activity (i.e., SIS3 + picrotoxin) induces proliferation of glioblastoma cells to levels significantly higher than those of SIS3-treatment alone (Fig. 6e,f). This effect in proliferation is not observed upon treatment with the Alk5 inhibitor A83 alone; however, the combination of A83 and picrotoxin also leads to increased proliferation compared to untreated cells (Fig. 6g,h), though to a lesser extent than upon SIS3 treatment. Noteworthy, TGFβ signalling inhibition in combination with picrotoxin treatment only promotes proliferation of the glioblastoma cells in co-culture with glutamatergic neurons, not when cultured in the absence of neurons (Supplementary Figure 6b). In line with our multi-omics data, these functional assays suggest that inhibition of SMAD3 and stimulation of neural activity additively cooperate to promote cell proliferation in glioblastoma cells in co-culture with glutamatergic neurons.

**Figure 6:**
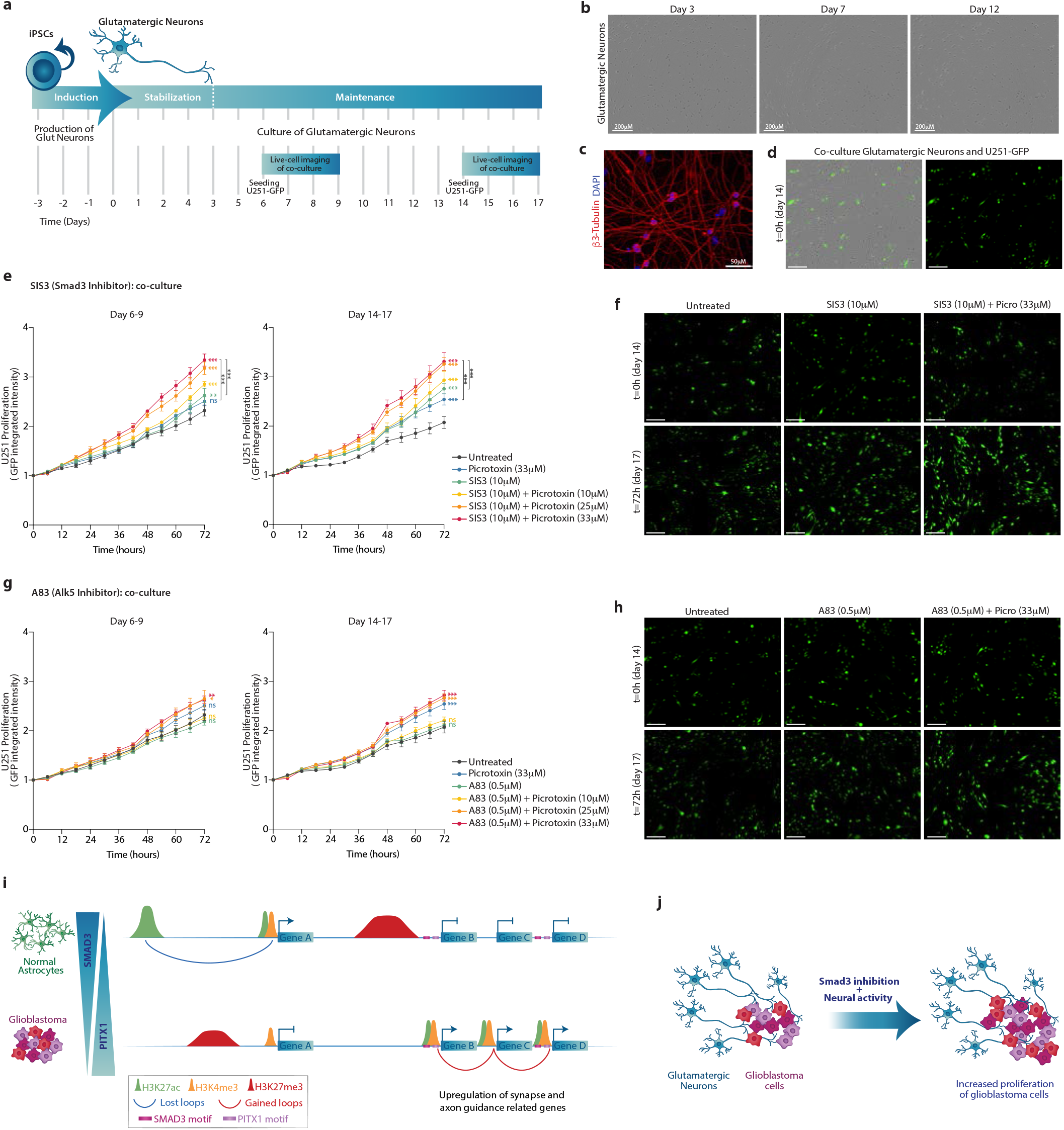
SMAD3 inhibition and neuronal activity cooperate to promote proliferation of glioblastoma cells in co-culture with glutamatergic neurons. a) Workflow to reprogram glutamatergic neurons from iPSCs, establish a co-culture with GFP-labelled U251 glioblastoma cells and live-cell image acquisition. b) Photomicrograph of glutamatergic neurons at days 3, 7 and 12. c) Glutamatergic neurons stained with an antibody against β3-Tubulin (red) (DAPI, blue). d) Photomicrographs of glutamatergic neurons and U251-GFP glioblastoma cells (green) in co-culture. e,g) Proliferation curves depicting the growth of U251 cells (measured as normalized GFP integrated intensity) in co-culture with glutamatergic neurons in the 72h after seeding at day 6 (left) or day 14 (right), while treated with either the SMAD3-specific inhibitor SIS3 (e) or the ALK5 inhibitor A83 (g), in combination with increasing concentrations of picrotoxin. (Multiple unpaired t-test, t=72h, (* p<0.01, ** p<0.005, *** p<0.000005, n=6-8=). f,h) Representative images from the live-cell imaging proliferation assay of U251-GFP in co-culture with glutamatergic neurons, in the presence of either SIS3 (f) or A83 (h). i) Scheme illustrating the regulatory and topological changes in the glioblastoma genome including loss of long-range interactions, gain of promoter-promoter loops, redistribution of active chromatin marks towards SMAD3 and PITX1 binding motifs and upregulation of genes related to synapse organization, axon guidance and axonogenesis. j) Illustration depicting that Smad3 inhibition and stimulation of neural activity cooperate to promote proliferation of glioblastoma cells in co-culture with glutamatergic neurons.

## Discussion

Up to now, few recent reports had integrated chromatin/epigenetic profiling and transcriptomics in glioblastoma^26–29^, and much effort had been put on identifying distinct molecular features of GB subtypes^23,24^. This is of great importance to identify clinically relevant subpopulations with the goal of improving the outcome, however it has not yet resulted into clinical benefits. Here, using a broad panel of patient-derived glioblastoma cell lines and normal human astrocytes as control, we identified changes in the promoter-enhancer interactome, chromatin accessibility and redistribution of histone marks that are present across all four GB expression subtypes (Fig. 6i). Such rewiring of the regulatory landscape and 3D organization of the glioblastoma genome orchestrates gene expression changes which underlie neurogliomal synaptic communication. This is manifested by changes in the expression of genes related to synapse organization, axon guidance and axonogenesis, as well as chromatin binding/remodeling.

Chromatin profiling revealed a preferential loss of long-range regulatory loops and reduction of the active mark H3K27ac at strong and weak enhancers in glioblastoma, together with overall activation of promoters, as evidenced by higher enrichment of active marks at poised and weak promoters and reduction of the repressive mark H3K27me3 at those sites. Motif analysis reveals a significant enrichment of SMAD3 and PITX1 sites within gained active and open regions that are located at the promoters of genes related to synapse organization, in particular glutamatergic synapses, axon guidance and axonogenesis. Among the common SMAD3/PITX1 targets, it is worth highlighting genes such as *SEMA5A* and *EPHB3*, classical axon guidance molecules^42,43^; the transcription factor *NKX2-2* involved in axon guidance^45^; *HCN2* and *KCNE4* encoding for gated channels; and *TNIK* involved in glutamatergic synaptic function^47^. Moreover, regulatory and topological alterations in the *SMAD3* and *PITX1* loci are accompanied by changes in the expression of both *SMAD3* and *PITX1* genes, which are inversely correlated and associated to lower survival rates in patients. Even if additional transcription factors (i.e., EBF1, THRB, EN1, HOXB13, HOXD10) could contribute by regulating distal genes involved in axonogenesis/axon guidance, our data altogether suggests that SMAD3 and PITX1 act as major direct regulators of a set of downstream target genes related to synapse organization, glutamatergic synapses and axon guidance in glioblastoma. Interestingly, SMAD3 and PITX1 motifs are also significantly enriched at the anchors of differential loops, raising the question of whether they have functions directly linked to chromatin organization. Previous reports have shown that SMAD3 interacts with CTCF in mammalian cells^49^, and Smad proteins bind CTCF sites in a CTCF-dependent manner in flies^48^. However, whether PITX1 interacts with CTCF or to what extent SMAD3 and PITX1 contribute to 3D chromatin organization are aspects that remain to be explored.

The recently discovered neurogliomal synapses provide glutamatergic synaptic input that drives tumour progression^31,32^, induces formation of tumour microtubes and speeds up tumour cell invasion by hijacking neuronal migration mechanisms^33^. Importantly, using co-cultures of GB cells and glutamatergic neurons, we functionally demonstrated that inhibition of TGFβ signalling, and specifically SMAD3, in combination with stimulation of neural activity, promotes the proliferation of glioblastoma cells. Cell type-specific transcription factors^50^ and epigenomes^51^ can modulate SMAD3 downstream targets and therefore orchestrate cell type-specific effects of TGF-β signalling, a pathway which in cancer has dual roles in the regulation of cell death and proliferation depending on the context^52^. This is noteworthy since glioblastoma cells can transition between states under selective pressure (e.g., in response to treatments), which may also result in different effects when comparing different *in vivo* and *in vitro* experimental set-ups. Even though SMAD3 inhibition might reduce cell viability of certain GB subtypes^53^, our data underlines the importance of the cellular context, i.e. contacts with neurons, where the combination of neural activity stimulation and SMAD3 inhibition have proven to additively cooperate to promote proliferation of glioblastoma cells (Fig. 6j). Considering these neural-cancer interactions will be pivotal to improve the prognosis of malignancies difficult to treat, such as glioblastoma. Our study thus provides details of the regulatory and topological alterations in glioblastoma and offers novel mechanistic insight into the gene regulatory networks that mediate the neurogliomal synaptic communication.

## Acknowledgements

We acknowledge the Biochemical Imaging Center at Umeå University and the National Microscopy Infrastructure, NMI (VR-RFI 2019-00217) for providing assistance in microscopy. Research in SR’s laboratory is supported by the Swedish Research Council (2019-01960), the Swedish Cancer Foundation (21 1720), Knut och Alice Wallenbergs Stiftelse (WCMM), the Kempe Foundation (SMK-1964.2) and the Cancer Research Foundation Norrland (AMP 19-977, LP 21-2290, AMP 22-1091). This study is dedicated to the memory of J.M. Remeseiro.

## Authors contributions

CC and IN designed, performed, analysed and interpreted most of the experiments with contributions from CV and AC-H. CV performed and analysed the live-cell imaging experiments. AH co-supervised some bioinformatic analysis, and contributed to data interpretation and manuscript editing. SR designed and supervised the study, secured funding, analysed and interpreted the data, and wrote the manuscript with input from the other authors.

These authors contributed equally: CC and IN.

Corresponding author: Silvia Remeseiro

---

**Supplementary Figure 1:**
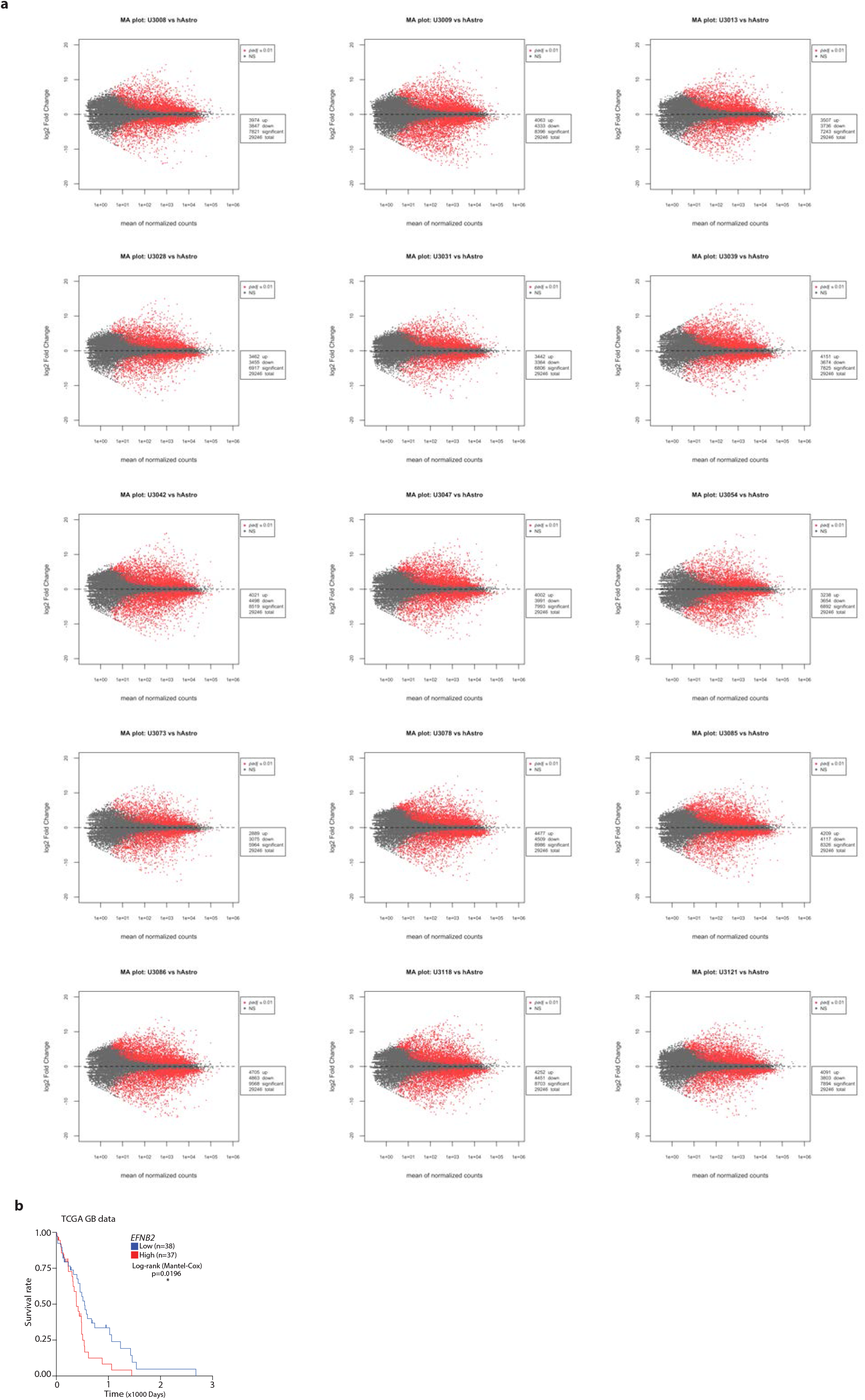
a) MA plots showing genes differentially expressed (DEGs, red dots) in each of the 15 glioblastoma lines *vs* normal astrocytes. b) Kaplan-Meier survival curves plotted for patients stratified by quartiles of *EFNB2* expression (TCGA data) (log-rank Mantel-Cox test).

**Supplementary Figure 2:**
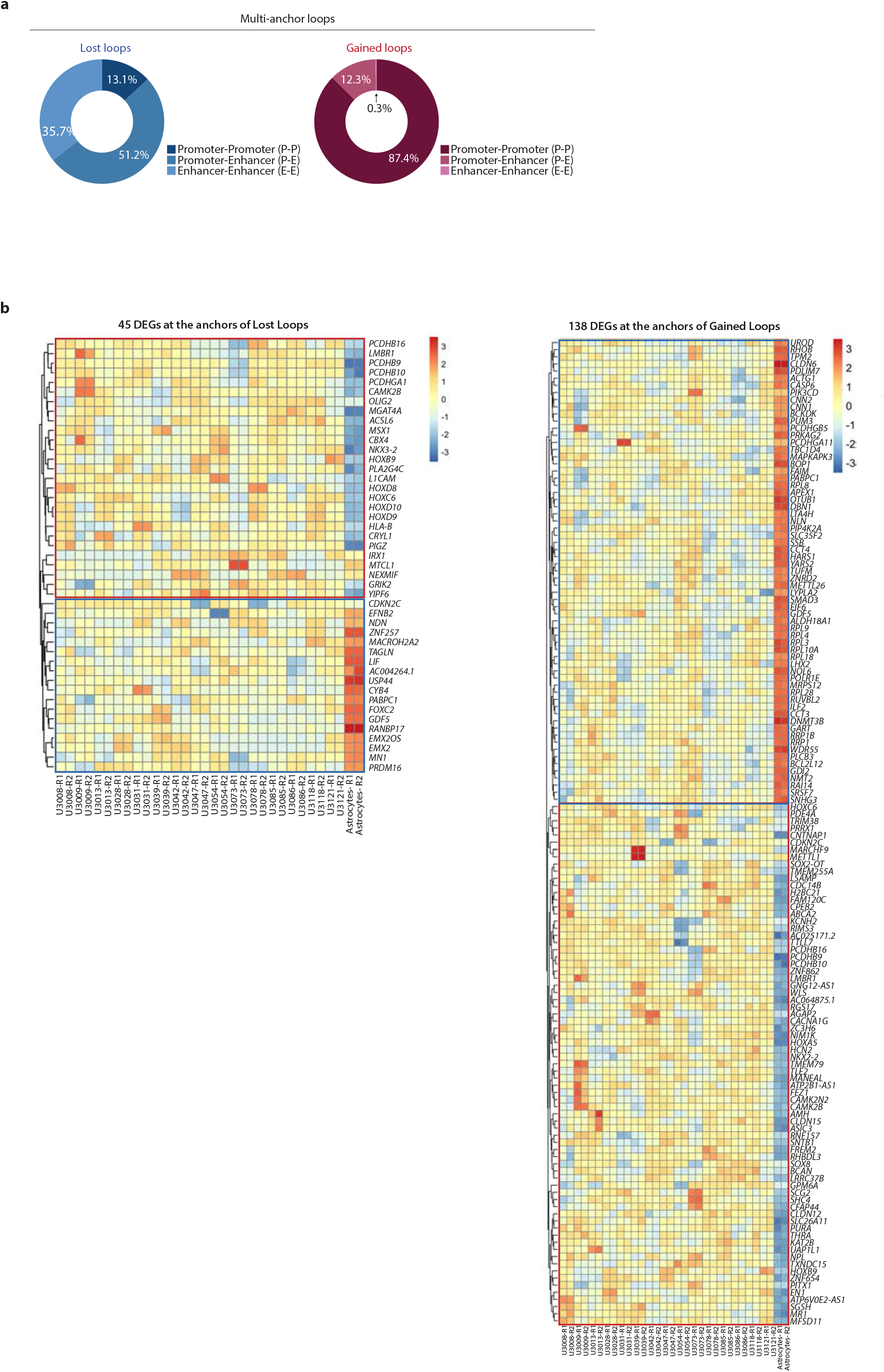
a) Annotation of lost and gained multi-anchor loops as promoter-promoter, promoter-enhancer or enhancer-enhancer loops. b) Expression of 45 DEGs located at the anchors of lost loops (left) and 138 DEGs at the anchors of gained loops (right), in all 15 glioblastoma lines and normal astrocytes as determined by RNA-seq (rlog normalized counts).

**Supplementary Figure 3:**
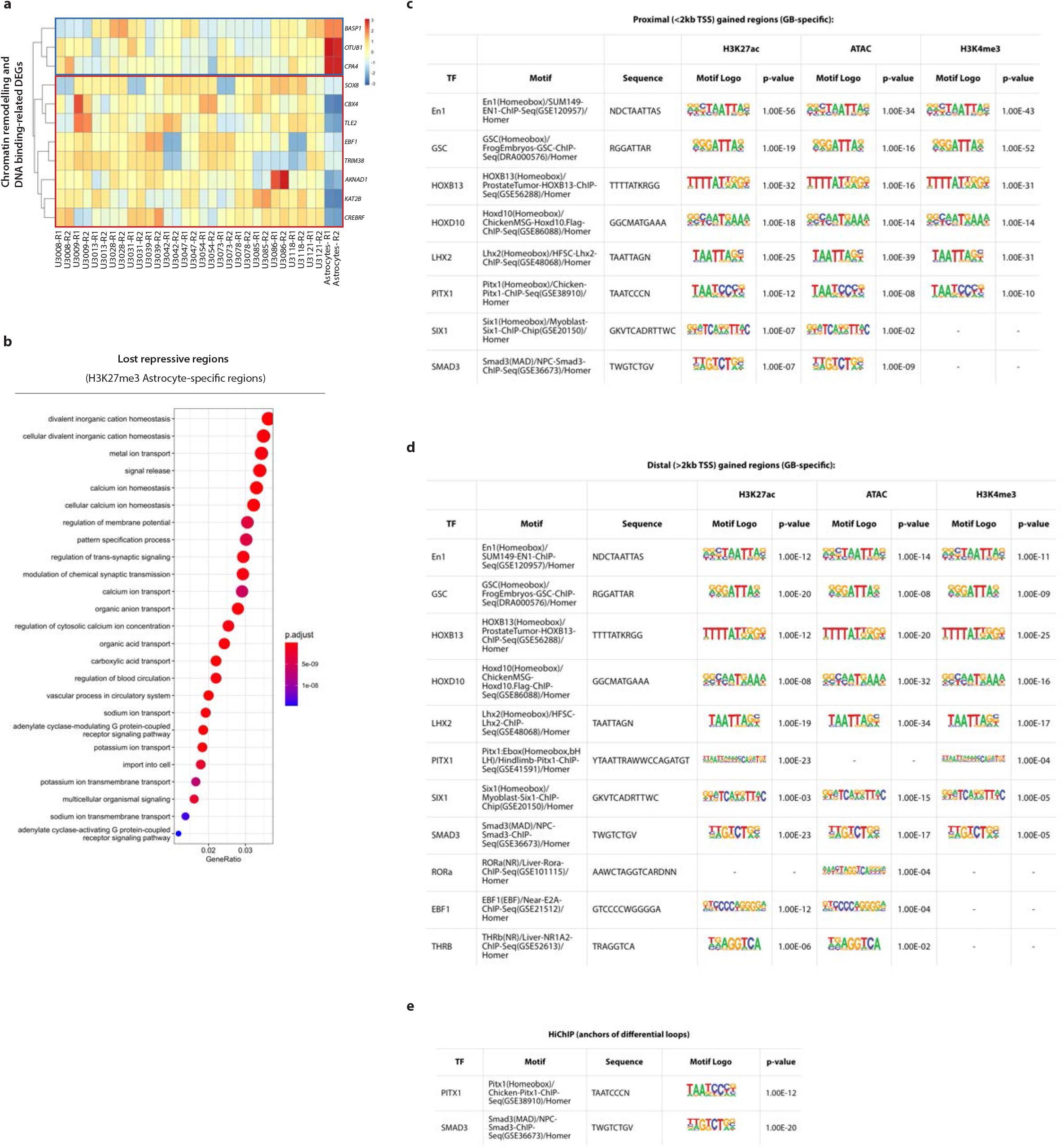
a) Expression of DEGs related to chromatin remodelling and DNA binding located <2kb from a gained active/open peak. b) Top 25 GO terms enriched for the genes proximal to the lost repressive regions in glioblastoma (peaks <2kb TSS). c,d) Summary of the TF binding motifs enriched at proximal (<2kb TSS) (c) or distal (>2kb TSS) (d) gained regions in glioblastoma for H3K27ac, ATAC and H3K4me3, including motif, sequence and p-values. e) Summary of the TF binding motifs enriched at the anchors of the differential loops, including motif, sequence and p-value.

**Supplementary Figure 4:**
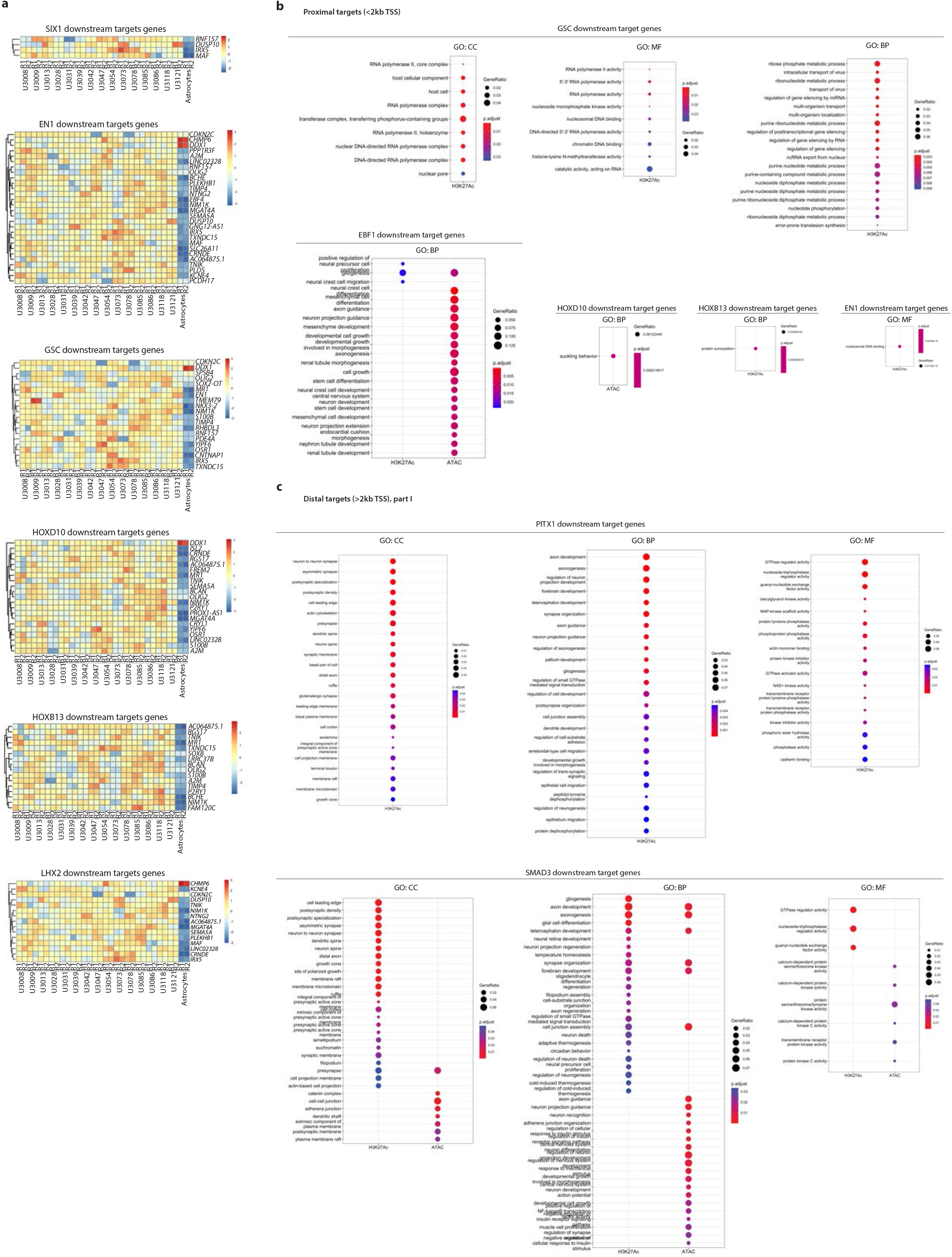
a) Expression of the SIX1, EN1, GSC, HOXD10, HOXB13 and LHX2 downstream target genes (i.e., motif enrichment at a differential peak <2kb TSS) that are DEGs in glioblastoma *vs* normal astrocytes. b) GO terms enriched for the GSC, EBF1, HOXD10, EN1 and HOXB13 proximal downstream target genes (<2kb TSS). c) GO terms enriched for the PITX1 and SMAD3 distal downstream target genes (>2kb TSS).

**Supplementary Figure 5:**
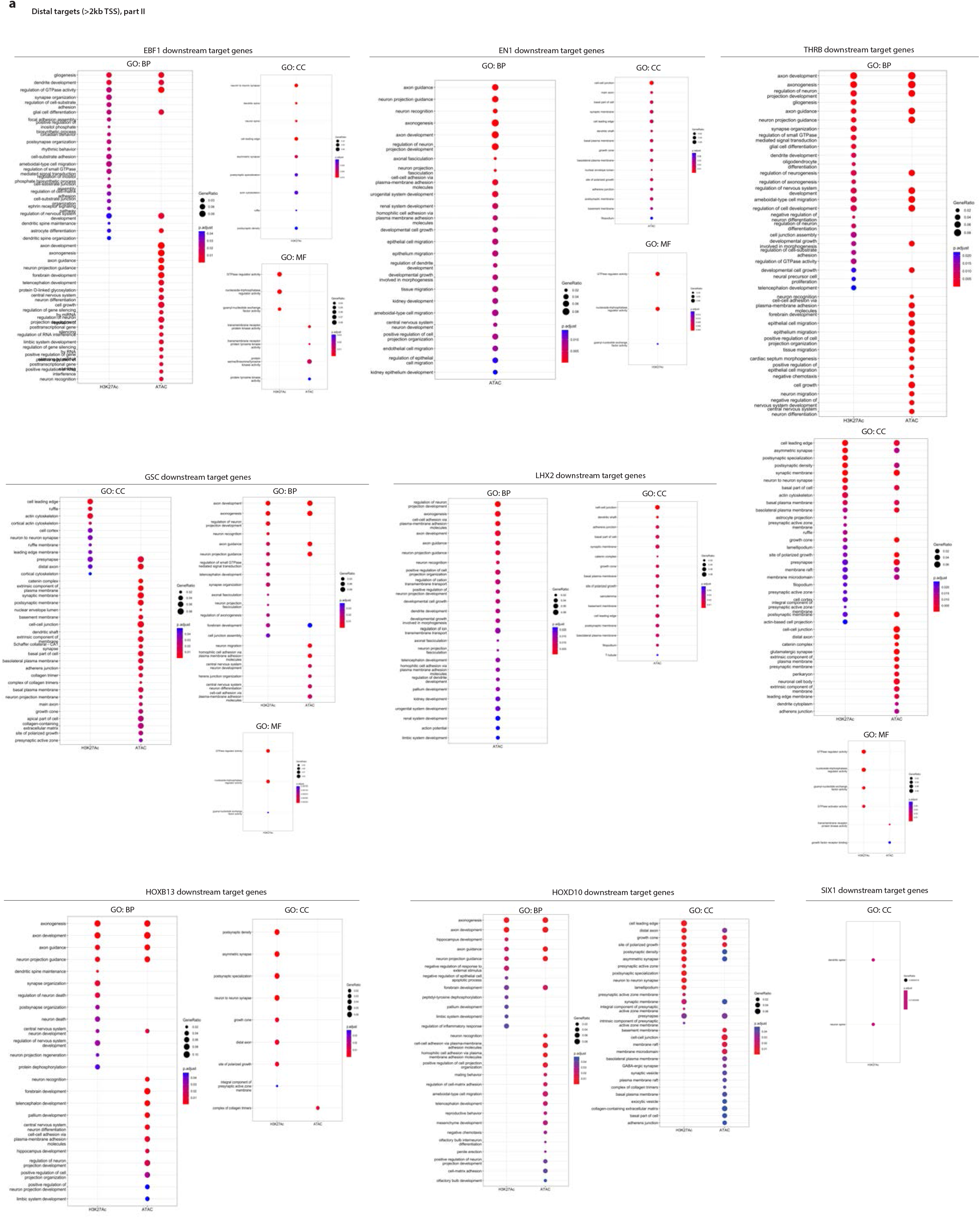
a) GO terms enriched for the EBF1, EN1, THRB, GSC, LHX2, HOXB13, HOXD10 and SIX1 distal downstream target genes (>2kb TSS).

**Supplementary Figure 6:**
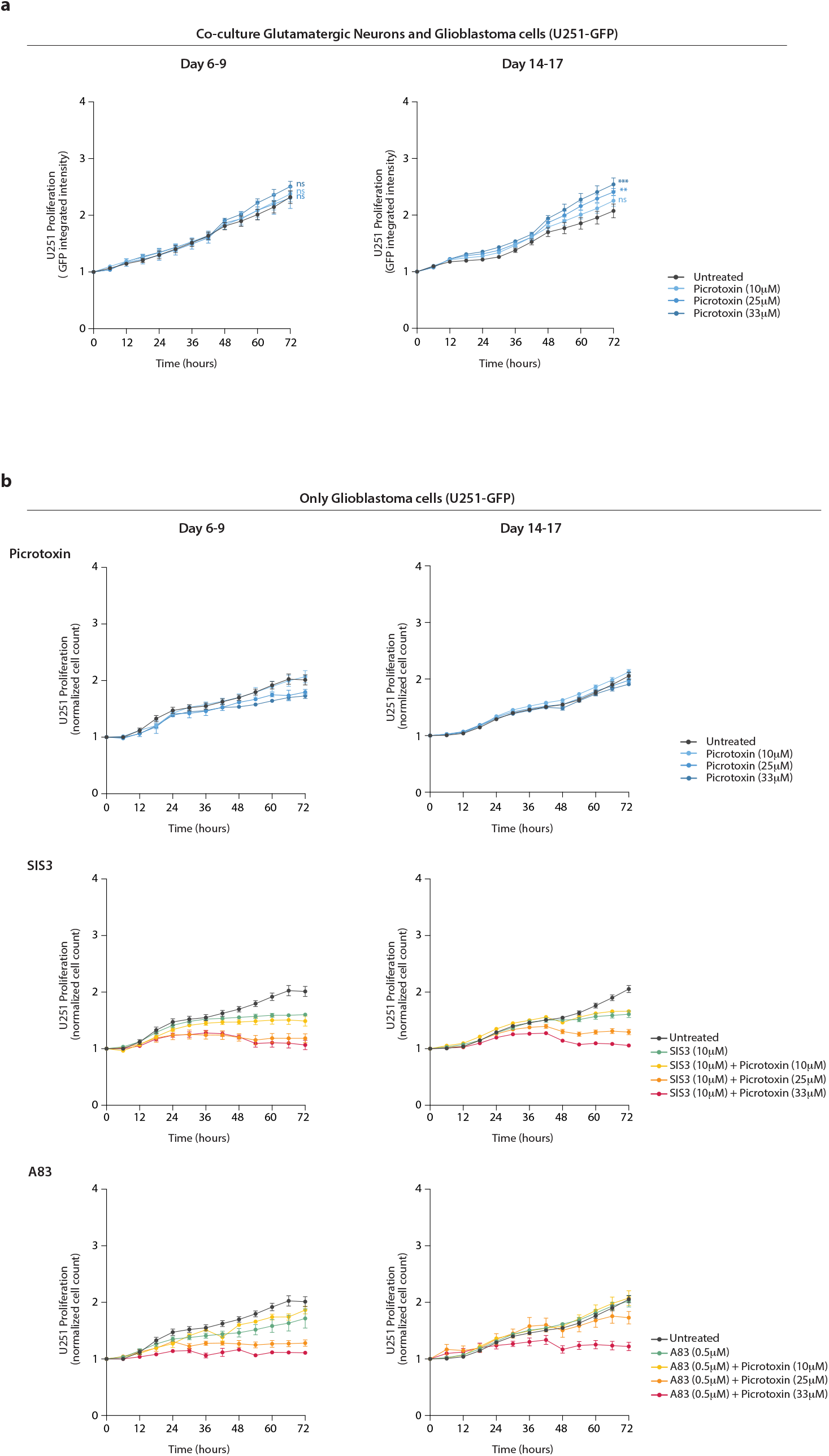
a) Proliferation curves depicting the growth of U251 cells (measured as normalized GFP integrated intensity) in co-culture with glutamatergic neurons in the 72h after seeding at day 6 (left) or day 14 (right) at different concentrations of picrotoxin. (Multiple unpaired t-test, t=72h, (* p<0.01, ** p<0.005, *** p<0.000005). (n=6-8). b) Proliferation curves of U251 cells in the absence of neurons (measured as normalized cell count) in the 72h after seeding at day 6 (left) or day 14 (right) and under the same drug treatments as used in Figure 6e,g.

## Methods

### Cell Culture

*Human glioblastoma cell lines* were derived from glioblastoma (GB) patient biopsies and obtained via the Human Glioblastoma Cell Culture (HGCC) resource^1^ (Uppsala University, Sweden). The 15 patient-derived glioblastoma lines represent the four GB expression subtypes (n=5 classical, n=5 mesenchymal, n=3 pro-neural and n=2 neural). Cells were seeded onto poly-ornithine/laminin-coated plates and grown in Feed Medium (1:1 ratio of DMEM/F12 Glutamax (Gibco) and Neurobasal medium (Gibco), supplemented with 1X B27 (Gibco), 1X N-2 Supplement (Gibco), 1% penicillin/ streptomycin (Gibco), 10 ng/ml EGF (Epithelial Growth Factor; PreproTech EC Ltd.) and 10 ng/ml FGF (Fibroblast Growth Factor; PeproTech EC Ltd).

*Normal Human Astrocytes* (Lonza, CC-2565) were grown in AGM Astrocyte Growth medium BulletKit (Lonza, CC-3186).

*U251 glioblastoma cells* (Sigma-Aldrich, #09063001) were grown in EMEM (EBSS) supplemented with 2mM Glutamine, 1% NEAA (Non-Essential Amino Acids), 1mM Sodium Pyruvate, 10% FBS (Fetal Bovine Serum) and 1% penicillin/streptomycin (all from Gibco). The GFP-labelled U251 line was established by lentiviral integration of the GFP reporter gene.

The *iPSC-derived glutamatergic neurons* (ioGlutamatergic neurons) were purchased from Abcam (ab259259), where human iPSCs were exposed to a 3-day induction protocol (day −3 to 0) and ioGlutamatergic neurons were cryopreserved. Upon thawing, 11 000 ioGlutamatergic neurons per well were seeded onto a PDL-Geltrex-coated 96-well plates and grown in Complete Glutamatergic Neuron Medium (CGNM) i.e., Neurobasal medium (Gibco) supplemented with 1X Glutamax (Gibco), 25μM 2-mercaptoethanol (Gibco), 1X B27 (Gibco), 10 ng/ml NT3 (R&D) and 5 ng/ml BDNF (R&D). During the stabilization phase, the CGNM was supplemented with 1 μg/ml doxycycline (Sigma) during 96h (day 0-4) and DAPT (Sigma) for 48h (day 2-4) for sustained induction. During the maintenance phase (day 4 onwards), the ioGlutamatergic neurons were grown in CGNM (without doxycycline and DAPT) with a half-medium change regime every 48 hours.

To establish the *co-culture of glutamatergic neurons and glioblastoma cells*, 5 000 U251-GFP glioblastoma cells were seeded onto the glutamatergic neurons either at day 6 or day 14 of culture. The co-cultures were treated with Picrotoxin (10, 25 or 33 μM, TOCRIS #1128), SIS3 (10 μM, Calbiochem #566405), A83 (0.5 μM, Sigma SML0788) or combinations of Picrotoxin and SIS3/A83. Live-cell imaging was performed on an IncuCyte S3 Live-Cell Analysis instrument (Sartorius) and proliferation of the GFP-labelled U251 cells was determined by measuring GFP integrated intensity using the Incucyte Base Analysis Software. Data points in Figure 6e,g and Supplementary Figure 6 correspond to n=6-8 replicates per condition and timepoint, for which 4 fields were imaged, and values are normalized to t=0. Significant differences in cell proliferation were assessed using unpaired t-test with correction for multiple testing at t=72h (* p<0.01, ** p<0.005, *** p<0.000005).

All cell lines were grown in a cell incubator at 37°C in a humidified atmosphere (95% humidity) with 5% CO2.

### RNA-seq

RNA-seq was performed in 15 glioblastoma cell lines and normal human astrocytes. Total RNA was extracted using RNeasy Plus Mini Kit (Qiagen, #74134) in technical duplicates for each cell line. Poly(A) RNA was purified using the NEBNext Poly(A) mRNA Magnetic Isolation Module (CAT #E7490L). RNA-seq libraries were prepared using the NEBNext® Ultra™ II RNA Library Prep Kit for Illumina (CAT# E7770L) and NEBNext® Multiplex Oligos for Illumina® (96 Unique Dual Index Primer Pairs) (CAT #E6440S) following the manufacturer’s instructions (8 amplification cycles). The RNA-seq libraries were sequenced on a NovaSeq 6000 Sequencing System (Illumina) obtaining in average ~53 million 150PE reads per library.

#### RNA-seq analysis

Fastq files were quality-checked with FastQC (https://www.bioinformatics.babraham.ac.uk/projects/fastqc/) and raw reads were mapped to the human genome (GRCh38/hg38) using STAR (2.7.6a)^2^. Genes with a minimum row sum of 10 reads were kept for further analysis. Normalization and differential expression analysis of each glioblastoma line *vs* the control normal astrocytes were performed using DESeq2 (p<0.01 and FDR<0.01), and the DEGs resulting from the 15 pair-wise comparisons were then intersected using the UpSet command from the ComplexHeatmap package, resulting in 497 differentially expressed genes (DEGs) across all 15 glioblastoma lines. Pheatmap package was used for clustering of differentially expressed genes in figures 1d, 4b, 4d, and supplementary figures 2b, 3a and 4a. Gene Ontology (GO) and KEGG enrichment analysis were performed using clusterProfiler (p<0.01 and FDR<0.01)^3^ (CC: cellular component, MF: molecular function, BP: biological process). RNA-seq data available at The Cancer Genome Atlas (TCGA) from both human Glioblastoma (GB, n=156) and Low-Grade Glioma (LGG, n=511) tissue samples were retrieved using the R packages TCGAbiolinks and RTCGAToolbox, together with normal tissue samples. We calculated Euclidean distances among the tumour samples based on the gene expression of our 497 DEGs and plotted the sample-to-sample distances as a heatmap. The differential expression analysis of either GB samples or LGG samples *versus* the respective normal control tissues was performed using DESeq2 (FDR<0.01). The UpSet command from the ComplexHeatmap package was used to intersect the 497 DEGs from our data with the DEGs resulting from the TCGA-GB *vs* Normal and TCGA-LGG *vs* Normal differential expression analysis, and the intersection was represented as an UpSet plot. Co-expression analysis for *SMAD3-PITX1* and *SMAD3-SIX1* gene pairs and Pearson correlation was calculated and plotted on R and ggplot2, using the rlog normalized counts for our RNA-seq data and the vsd log2 normalized counts for the TCGA data.

### ChIP-seq

ChIP-seq was performed in 15 glioblastoma cell lines and normal human astrocytes as described before^4,5^ with some modifications. Briefly, cells were fixed on the plate by adding formaldehyde directly to the medium (final concentration 1% formaldehyde) for 15 minutes at room temperature while rotating. The crosslinking reaction was quenched by adding Glycine (final concentration 125mM Glycine) for 5 min, and fixed cells were scraped off and harvested in 1X cold PBS containing protease inhibitors. Cells were then resuspended in lysis buffer (3-6×10^6^ cells/ml) and sonicated in a Covaris E220 instrument (shearing time 12min, PIP 140, duty factor 5, 200 cycles per burst), to achieve a fragment size ranging from 200bp to 700bp. Chromatin immunoprecipitation was performed with antibodies against H3K27ac (ab4729), H3K4me3 (ab8580) and H3K27me3 (ab192985), and using Dynabeads™ M-280 Sheep Anti-Rabbit IgG (Invitrogen 11203D). H3K27ac and H3K27me3 ChIP–seq libraries were prepared using the NEBNext Ultra II DNA Library Prep Kit for Illumina (E7645L) and NEBNext® Multiplex Oligos for Illumina® (E6440S). The H3K4me3 library was prepared using Accel NGS 2S Plus DNA Library Prep (#21024, Swift Biosciences) and indexing Kit (#26596, Swift Biosciences), since it was processed and sequenced together with the H3K4me3-HiChIP. ~10ng of immunoprecipitated chromatin (as quantitated by the Qubit fluorometer) alongside the corresponding inputs were amplified for 8 cycles and further processed according to the guidelines of the library prep kits. SPRI Select beads (Beckman Coulter) were used for clean-up and size selection. The ChIP-seq libraries were sequenced on a NovaSeq 6000 Sequencing System (Illumina) obtaining in average ~51 million 150PE reads per library.

#### ChIP-seq analysis

Fastq files were quality-checked with FastQC (https://www.bioinformatics.babraham.ac.uk/projects/fastqc/) and raw reads were mapped to the human genome (GRCh38/hg38) using bowtie2 (2.4.1)^6^ (--threads 4 --very-sensitive). Peak calling was performed by MACS2^7^ (options: --broad -g hs -B -q 0.05 -f BAMPE) using the corresponding input track as control (i.e. astrocytes *vs* astrocyte input, and each of the glioblastoma cells *vs* the corresponding input depending on the subtype i.e., CL-input, MS-input, PN-input or NL-input). ChIP-seq signal was plotted as heatmaps using plotHeatmap from deepTools. Bigwig files were visualized in the UCSC genome browser (https://genome.ucsc.edu/). Density plots displaying the signal around TSSs or defined chromatin states were performed with plotProfile upon calculation of enrichment using computeMatrix. Bedtools multiinter tool was used to determine the regions that were lost or gained in glioblastoma for each of the histone marks. Lost regions were defined as astrocyte-specific regions (i.e. present in astrocytes and absent in all 15 glioblastoma lines), while gained regions were defined as glioblastoma-specific (i.e. absent in astrocytes and present in at least 10 out of 15 glioblastoma lines). Genomic annotations of peaks as well as lost and gained regions were obtained using BSgenome.Hsapiens.UCSC.hg38, ChIPpeakAnno, ChIPseeker^8^ & EnsDb.Hsapiens.v86 in R. Gene Ontology analysis of genes proximal (<2kb TSS) or distal (>2kb TSS) to the gained/lost regions were performed using clusterProfiler^3^ (CC: cellular component, MF: molecular function, BP: biological process). Search for TFBS (transcription factor binding sites) motifs was conducted using HOMER (v4.11) tool findMotifsGenome.pl. For each histone mark, motif analysis was performed within the differential regions located either proximally (<2kb) or distally (>2kb) to TSSs. Significantly enriched TF motifs (p<0.01) were intersected with the 497 DEGs to identify transcription factors that were differentially expressed and whose motifs were enriched at the differential histone regions. Distances between SMAD3 and PITX1 motifs at the promoters of the 33 common SMAD3/PITX1 target genes were calculated within the differential regions (i.e., gained H3K27ac and ATAC peaks) located <2kb of TSS, and using as background model the common peaks at promoters genome-wide. Discovery of chromatin states in normal astrocytes was performed with ChromHMM^9^ using the H3K27ac, H3K4me3, H3K27me3, ATAC-seq and RNA-seq datasets as input and setting 8 state emissions.

### ATAC-seq

ATAC-seq was performed in 15 glioblastoma cell lines and normal human astrocytes (50 000 cells/sample) as previously described^10,11^. Briefly, DNA tagmentation was performed 30 minutes at 37°C using the Illumina Tagment DNA TDE1 Enzyme and Buffer Kits (#20034197). The reaction was purified using a MinElute Purification Kit (Qiagen #28004) and fragmentation was assessed via Bioanalyzer High Sensitivity DNA Analysis (Agilent # 5067-4626). 5μl of tagmented DNA per library were amplified for 13 cycles using NEBNext High-Fidelity 2x PCR Master Mix (M0541S) and custom oligonucleotides (for oligo sequences see Supplementary Table S3). The ATAC libraries were sequenced on a NovaSeq 6000 Sequencing System (Illumina) aiming ~55 million 150PE reads per library.

#### ATAC-seq Analysis

ATAC-seq analysis was performed according to the ENCODE ATAC-seq Processing Pipeline with some modifications. Fastq files were quality-checked with FastQC (https://www.bioinformatics.babraham.ac.uk/projects/fastqc/). The pipeline for further processing included trimming with cutadapt to remove the Nextera adaptor sequence CTGTCTCTTATACACATCT, mapping to the human genome (GRCh38/hg38) using bowtie2^6^ (--k 2, --threads 8, --local, --maxins 2000), removing duplicates with MarkDuplicates from Picard toolbox and filtering with samtools to keep high quality and uniquely aligned read pairs. Peak calling was performed using MACS2^7^ (--broad -q 0.05 - -shift 100 --extSize 200 against baseline). Genomic annotations of peaks were obtained using ChIPseeker^8^; the ATAC-seq signal was plotted as heatmaps using plotHeatmap from deepTools, and bigwig files were visualized in the UCSC genome browser (https://genome.ucsc.edu/). Density plots displaying the signal around defined chromatin states were performed with plotProfile upon calculation of enrichment using computeMatrix. Bedtools multiinter tool was used to determine the ATAC regions that were lost or gained in glioblastoma. Lost regions were defined as astrocyte-specific regions (i.e. present in astrocytes and absent in all 15 glioblastoma lines), while gained regions were defined as glioblastoma-specific (i.e. absent in astrocytes and present in at least 10 out of 15 glioblastoma lines). Lost and gained ATAC regions were then annotated using BSgenome.Hsapiens.UCSC.hg38, ChIPpeakAnno, ChIPseeker^8^ & EnsDb.Hsapiens.v86 in R. Gene Ontology analysis of genes proximal (<2kb TSS) or distal (>2kb TSS) to the gained/lost ATAC regions were performed using clusterProfiler (CC: cellular component, MF: molecular function, BP: biological process). Search for TFBS (transcription factor binding sites) motifs was conducted using HOMER (v4.11) tool findMotifsGenome.pl. Motif analysis was performed within the differential ATAC regions located either proximally (<2kb) or distally (>2kb) to TSSs. Significantly enriched TF motifs (p<0.01) were intersected with the 497 DEGs to identify transcription factors that were differentially expressed and whose motifs were enriched at the differential ATAC regions.

### HiChIP

HiChIP was performed in 15 glioblastoma cell lines and normal human astrocytes using the Arima-HiC+ Kit (A101020) and following the guidelines in the Arima-HiChIP user guide for mammalian cells. 9-15×10^6^ cells per line were used to obtain at least 12-15μg of input DNA for HiChIP. Cells were fixed in 1% formaldehyde for 15 min at room temperature while rotating, and crosslinking reaction was quenched by adding Glycine (final concentration 125mM Glycine) for 5 min. Subsequent steps included digestion of crosslinked chromatin with restriction enzymes, end-filling with biotinylated nucleotides and ligation. Proximally ligated chromatin was then sheared on a Covaris E22O instrument (shearing time 5min, PIP 105, duty factor 5, 200 cycles), to achieve a fragment size ranging from 200bp to 800bp. Chromatin immunoprecipitation was performed with an antibody against H3K4me3 (ab8580). After biotin enrichment and adapters ligation, immunoprecipitated DNA was subjected to PCR amplification (8-11 cycles) using Accel-NGS 2S Plus DNA Library Kit (#21024, Swift Biosciences) and indexing Kit (#26696, Swift Biosciences), according to the Arima-HiChIP Library Prep user guide. Quality controls for chromatin digestion, ligation, shearing and library preparation were assessed via Bioanalyzer High Sensitivity DNA Analysis (Agilent # 5067-4626) and passed prior to sequencing. The HiChIP libraries were sequenced on a NovaSeq 6000 Sequencing System (Illumina) aiming ~100 million 150PE reads per library.

#### HiChIP analysis

HiChIP analysis was performed as described before^12^ using HiC-Pro and Hichipper. Fastq files were quality-checked with FastQC (https://www.bioinformatics.babraham.ac.uk/projects/fastqc/). Mapping to the human genome (GRCh38/hg38) and retrieval of valid interacting fragments was performed using the HiC-Pro software v3.1.0 and setting ligation sites as GATCGATC, GANTGATC, GANTANTC, GATCANTC. Valid loops were identified using Hichipper v.0.7.3. Significant loops were determined using diffloop (1.20.0)^13^ in R filtering for a minimum normalized read pair (loop count ≥ 2), FDR<0.01 and loop length ≥ 5000 bp. HiChIP samples passed quality control if ≥ 4000 significant loops were detected. To identify the differential loops, we defined lost and gained loops in glioblastoma as astrocyte-specific and GB-specific loops, respectively. The criteria were set such as lost loops are those present in normal astrocytes and absent in all glioblastoma lines{i.e., ∃ astrocytes (counts ≥2) & ∄ 14/14 GB (counts=0)}, while gained loops are absent in normal astrocytes and present in at least 8 out of the 14 glioblastoma lines{i.e., ∄ astrocytes (counts=0) & ∃ 8/14 GB (counts≥2)}. Annotation of the significant and differential loops were done using GenomicInteractions package in R (1.26.0). Loops were categorized as promoter-promoter (P-P), promoter-enhancer (P-E) or enhancer-enhancer (E-E) setting the criteria for promoter regions <2kb TSS and considering E-E as distal-distal regions. Differences in loop length (bp) between lost and gained loops were assessed using a t-test with Benjamini-Hochberg correction. Among the differential loops, multi-anchor loops were identified as loops that share anchors with other loop(s) i.e., one anchor is utilized by two or more loops. Search for TFBS (transcription factor binding sites) motifs was conducted using HOMER (v4.11) tool findMotifsGenome.pl. Motif analysis was performed within the anchors of the differential lost and gained loops. Significantly enriched TF motifs (p<0.01) were intersected with the 497 DEGs to identify transcription factors that were differentially expressed and whose motifs were enriched at the anchors of the differential loops.

### Statistical analysis

Statistical tests used are indicated in the corresponding Methods sections and in figure legends. Differences in loop length (bp) between lost and gained loops were assessed using a t-test with Benjamini-Hochberg correction. Kaplan-Meier survival curves were plotted for patients stratified by quartiles of expression and tested using a log-rank Mantel-Cox test. Differences in SMAD3/PITX1 motif distance were assessed using a Wilcoxon test. Differences in cell proliferation were determined using unpaired t-test with correction for multiple testing (* p<0.01, ** p>0.005, *** p<0.000005).

### Bioinformatic analyses and graphics

Most statistical analysis related to RNA-seq, ChIP-seq, ATAC-seq and HiCHIP were performed within the R environment (v.4.0.0-v.4.1.2) using basic built-in functions and publicly available packages dplyr, plyr, reshape2, ggmisc, ggpubr (R-CRAN). DESeq2, ComplexHeatmap, ClusterProfiler, BSgenome.Hsapiens.UCSC.hg38, ChIPpeakAnno, ChIPseeker and EnsDb.Hsapiens.v86 are open source tools available via Bioconductor (https://www.bioconductor.org/). Data in figures 1d, 2c, 2e, 2h, 5c, 6e, 6g and supplementary figures 2a and 6a,b were plotted using GraphPad Prism 9. The Circos Plot in figure 3a was generated using RCircos (R-CRAN package: 1.2.0). All other plots were created using basic R graphical interface and ggplot2. Genomic snapshots were downloaded from UCSC Genome Browser visualizations (https://genome.ucsc.edu/).

## Data availability

The RNA-seq, ChIP-seq, ATAC-seq and HiChIP datasets generated in this study have been deposited in GEO (Gene Expression Omnibus) under the accession number GSE217349.

